# Cdk1 and PP2A constitute a molecular switch controlling orderly degradation of atypical E2Fs

**DOI:** 10.1101/2025.02.23.639703

**Authors:** Sapir Nachum-Raines, Noy Gamliel, Danit Wasserman, Nasrin Qassem, Inbal Sher, Julia Guez-Haddad, Michael J Emanuele, Jordan H Chill, Amit Tzur

**Author notes:** Corresponding author Electronic address.

## Abstract

Dynamic oscillations in the phosphorylation and ubiquitination of key proliferative regulators are defining features of the eukaryotic cell cycle. Resetting the cell cycle at the mitosis-to-G1 transition requires activation of the E3 ubiquitin ligase Anaphase-Promoting Complex/Cyclosome (APC/C), which ensures cell cycle irreversibility by targeting dozens of substrates for degradation, safeguarding genome integrity. However, the overall coupling of substrate phosphorylation with target recognition and degradation by the APC/C remains relatively unexplored. As a paradigm for further defining these rules, we focused on E2F7 and E2F8 – atypical E2F-family proteins that coordinate cell cycle gene expression by restraining the pro-proliferative transcriptional activity of E2F1. Leveraging complementary cell and cell-free systems, we demonstrate that flexible domains in the amino-termini of E2F7 and E2F8 contain APC/C recognition motifs adjacent to critical Thr residues, whose phosphorylation by Cdk1 is rate limiting for degradation. The removal of this phosphorylation by PP2A phosphatase serves as a molecular switch, coupling the degradation of E2F7 and E2F8 to the G1 phase, coinciding with the rise of E2F1. Collectively, these findings highlight a critical role for Cdk1-PP2A signaling in controlling the orderly degradation of APC/C substrates, ensuring precisely timed assembly of the transcriptional infrastructure that coordinates cell cycle commitment and progression.

## Introduction

The cell cycle is controlled through fluctuations in the abundance and/or activity of proteins that perform specific functions at distinct stages, ensuring the accurate duplication and segregation of the genome [1]. These cyclic changes are controlled primarily at two levels: transcriptional regulation of gene expression and post-translational modifications, including ubiquitin-mediated proteolysis [2], which is crucial for driving the cell cycle in a unidirectional manner [3, 4].

The temporal regulation of proteins in cycling cells is additionally regulated by phosphorylation and dephosphorylation via kinases and phosphatases, whose activity, in turn, depends on regulatory proteins that oscillate throughout the cell cycle. For example, the key mitotic kinase Cdk1 is activated by cyclins A and B, whose accumulation and degradation are tightly controlled by the cell cycle clock at both transcriptional and proteolytic levels [1]. This intricate network of feedback mechanisms ensures the orderly progression of the cell cycle.

At the transcriptional level, the E2F family of transcription factors plays a pivotal role in coordinating cell cycle-dependent gene expression [5, 6]. The activator E2Fs (E2F1, E2F2, and E2F3a) promote the expression of genes required for the transition from the G1 phase of the cell cycle into DNA synthesis (S) phase. E2F7 and E2F8 (henceforth collectively referred to as E2F7/8) introduce an additional layer of regulation; these E2F members, also known as atypical E2Fs, are expressed during the S and G2 phases, where they suppress E2F-dependent transcription, modulating cell cycle control while also regulating E2F1-mediated DNA damage responses and apoptosis [7, 8]. As target genes of E2F1, E2F7/8 are effectively expressed as part of the cell cycle transcriptional program that they ultimately inactivate.

E2F1 activity commences at late G1 and rises sharply at the G1/S transition through an autocatalytic cycle [5]. Despite their transcriptional activation, atypical E2Fs do not accumulate until later in the S phase due to post-translational regulation; we and others have shown that both E2F members are targeted for degradation in G1 phase by the E3 ubiquitin ligase <u>A</u>naphase-<u>p</u>romoting <u>c</u>omplex/<u>C</u>yclosome (APC/C) [9, 10].

APC/C-mediated ubiquitination depends on two co-activators that confer substrate specificity in a phase-dependent manner—Cdc20 during mitosis and Cdh1 during cytokinesis and G1 [11]. Importantly, APC/C activity in G1 is progressively inhibited and ultimately terminated by E2F1 target gene products. This negative feedback loop is essential for ensuring the timely G1-to-S phase transition [11]. Furthermore, in G2 phase, E2F7/8 and E2F1-3a are regulated by the SCF E3 ligase through the substrate receptor Cyclin F [12–15]. Thus, the transcriptional activity of the E2F1-E2F7/8 circuitry is precisely timed and balanced by the ubiquitin-proteasome system throughout the cell cycle.

Timely activation of Cdc20-bound APC/C (APC/C^Cdc20^) and the transition to Cdh1-bound APC/C (APC/C^Cdh1^) during mitotic exit are regulated by the counteractive interplay between Cdk1 and protein phosphatase 2A (PP2A) along with other cell cycle kinases and phosphatases, ensuring orderly mitotic progression and exit [16–19]. Moreover, SCF-mediated ubiquitination is fundamentally dependent on the phosphorylation status of its substrates [20], further highlighting the coupling between (de)phosphorylation and the degradation of cell cycle regulators.

E2F7/8 contain multiple Cdk1 consensus sites clustered near the KEN-box motif [9, 14], a canonical APC/C recognition signal [21]. We have previously shown that E2F8 is phosphorylated by Cdk1 in mitosis and undergoes dephosphorylation at mitotic exit, mediated by an unidentified phosphatase [14]. However, it remains unclear whether and how E2F7/8 degradation is regulated, either directly or indirectly, by cell cycle kinases or phosphatases.

Understanding how E2F7/8 are regulated is important for defining their function in both homeostatic cell cycle progression and pathological conditions such as cancer [5, 6]. In this study, we aim to address this question using both conventional and newly developed cell-free approaches. These semi-physiological in vitro setups are optimal for quantitative, direct, and time-specific assays, overcoming uncertainties associated with long-term cellular manipulations [22–24]. We show that the orderly degradation of E2F7/8 is regulated by a phosphorylation switch mediated by the counteracting activities of Cdk1 and PP2A, highlighting a newly identified feedback mechanism that coordinates the transcriptional activity of the E2F1-E2F7/8 network with the cell cycle clock.

## Results

### Characterization of disorder in the N-terminal segments of E2F7 and E2F8

E2F7/8 are the largest members of the E2F family, consisting of 911 and 867 amino acid residues, respectively, which makes them challenging to characterize structurally. We have previously shown that the amino-terminal 80 amino acids of E2F8 are targeted by both Cdk1 and APC/C in a cell-cycle controlled manner [14]. These findings motivated us to investigate the structural properties of this E2F8 fragment (N80-E2F8) and the corresponding homologous region in E2F7, comprising residues 1-100 (N100-E2F7). Both fragments possess a KEN motif and a cluster of Cdk1 consensus sites **(Figure 1a)**. Following an unmatched 25-residue segment in the longer E2F7, the two sequences exhibit 47% identity and 59% similarity, which is strongest in the first 20 residues (63% similarity) and the last 40 residues (74%). Notably, this similarity correlates with the location of canonical Cdk1 consensus sites **(Figure 1a)**. Based on the IUPRED disorder prediction engine [25], both sequences are relatively disordered, with 71% of E2F7 and 100% of E2F8 scoring above the 0.5 disorder probability level **(Figure 1b)**.

**Figure 1.**
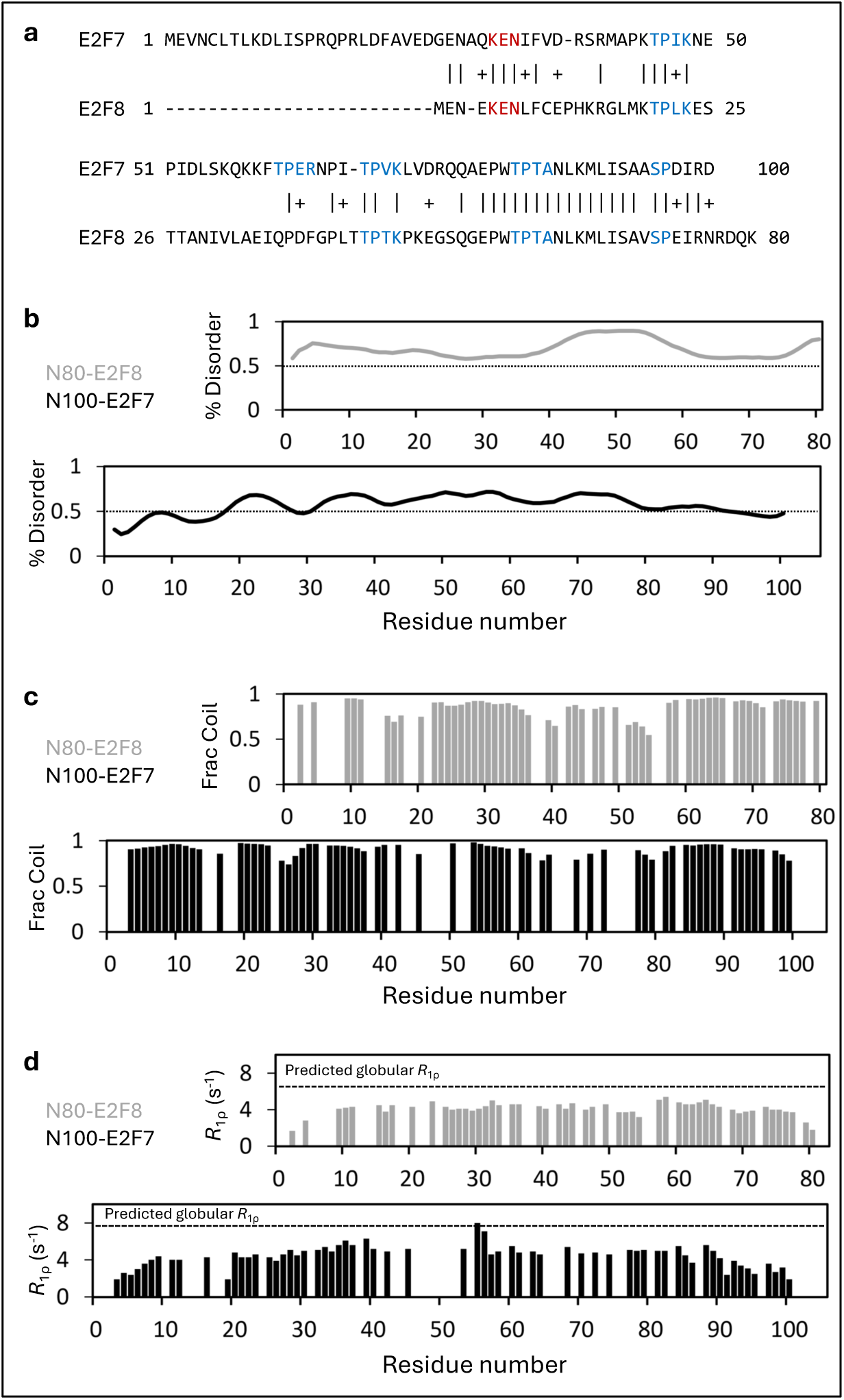
Characterization of disorder in N-terminal E2F7 and E2F8 fragments. **(a)** BLAST alignment of N100-E2F7 and N80-E2F8 sequences. KEN boxes and Cdk1 consensus sites are highlighted in red and blue, respectively. **(b)** IUPRED3 sequence-based prediction of disorder in the two sequences, with the 50% disorder threshold emphasized. **(c)** Delta2D NMR-based prediction of the fraction of random-coil (Frac coil) conformation for the two sequences. Missing data are due to insufficient resolution and/or signal-to-noise ratios, leading to ambiguity in chemical shift assignments. **(d)** NMR R1ρ relaxation rates at 16.4 T and 278 K for the two proteins. Expected rates for a globular protein of the same size are shown for comparison, indicating that both proteins are unfolded and disordered.

To experimentally verify the disordered nature of these segments we recombinantly expressed and purified ^15^N,^13^C-labeled N100-E2F7 and N80-E2F8 from *E. coli* grown in isotopically-labeled cultures. Both fingerprint two-dimensional (2D) ^1^H,^15^N-HSQC spectra exhibited low spectral dispersion characteristic of disordered protein segments. This constraint, together with the partial repetition within the E2F7/8 sequences, caused the assignment of their chemical shifts to be challenging. A combination of 3D-NMR spectra correlating ^1^H/^15^N/^13^C resonances along the E2F7/8 backbones provided reasonable assignment levels. Using the delta2D platform for NMR-based prediction of secondary structure [26], we translated backbone chemical shifts for 57 of the E2F7^1-100^ 90 non-proline residues (63%) and 51 of E2F8^1-80^ 72 non-proline residues (71%) into probabilities for finding the two sequences in random coil conformation. As shown in **Figure 1c**, the chemical shifts in both proteins are consistent with a mostly unfolded random coil conformation, averaging 91% for E2F7 and 86% for E2F8.

Further verification of the unstructured nature of these segments was provided by measuring transverse rotating-frame ^15^N auto-relaxation (^15^N-*R*_1ρ_) along the polypeptide backbones **(Figure 1D)**. Both fragments exhibited ^15^N-*R*_1ρ_ profiles typical of random-walk polymers, showing extremely low rates at the polypeptide termini (3-4 residues on each side) indicating enhanced flexibility, and a relatively flat profile of higher rates in the polypeptide central segment. ^15^N-*R*_1ρ_ rates in these central regions were (average ± STD) 4.9 ± 0.7 and 4.3 ± 0.4 s^−1^ for E2F7 and E2F8, respectively. For comparison, the expected relaxation rate for structured proteins of similar size would be 7.7 and 6.5 s^−1^ for E2F7 and E2F8, respectively [27], under our measurement conditions of 16.4 T and 278 K. Thus, the N-terminal segments of E2F7/8 share significant sequence similarity, and both proteins adopt a disordered and mostly unfolded conformation, based on predictive modeling and NMR. Interestingly, regulatory domains of other APC/C substrates are characterized by disordered conformation **(Figure S1** and Refs [28–30]).

### The E2F7 N-terminal domain is resistant to APC/C-mediated degradation

We have previously shown that N80-E2F8 can be targeted for ubiquitination and degradation by APC/C^Cdh1^ in a cell-free extract system that recapitulates G1-phase (G1 extracts) [14]. We took advantage of this biochemical system to investigate functional elements modulating E2F7 dynamics. The KEN motifs in N100-E2F7 and N80-E2F8 have been shown to mediate APC/C-dependent degradation of both full-length proteins [9, 14]. Unlike N80-E2F8, the degradation of the N-terminal fragment of E2F7 in G1 extracts has yet to be tested. To address this, the system’s potency and specificity were first validated using N80-E2F8 as a positive control substrate and a recombinant dominant-negative (DN) variant of the APC/C E2 enzyme UbcH10 (UbcH10DN), which blocks APC/C-specific activity [10]. Surprisingly, N100-E2F7 remained stable in G1 extracts **(Figure 2a)**, as did an extended fragment containing an additional 20 residues **(Figure S2)**.

**Figure 2.**
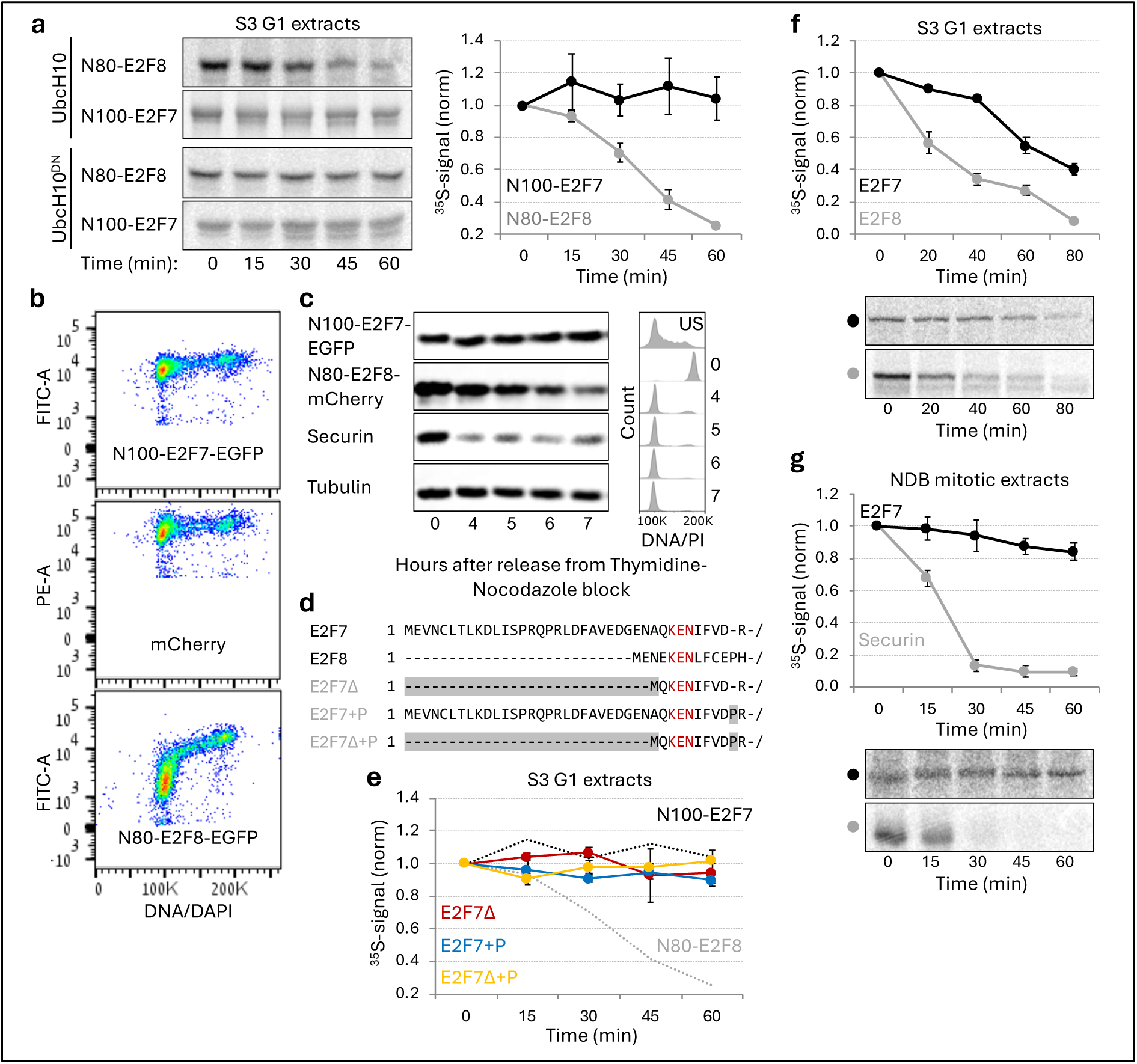
KEN-box-containing N100-E2F7 lacks degron properties. **(a)** Time-dependent degradation of N80-E2F8 and N100-E2F7 fragments in G1 extracts. Radiolabeled in vitro translated (IVT) products were incubated in HeLa S3 (S3) G1 extracts. Recombinant UbcH10 or UbcH10^DN^ were added to facilitate or block APC/C-specific activity, respectively. Representative source data are shown (left). Standard error (SEM) and normalized (norm; t_0_ = 1) mean ^35^S signals (right) representing N80-E2F8 and N100-E2F7 levels are plotted (n = 3). **(b)** Differential oscillation of E2F7/8 fragments in cycling HeLa cells. Bivariate plots present flow cytometry analysis of cells stably expressing N100-E2F7-EGFP, mCherry, or N80-E2F8-EGFP. Cells were fixed and stained with DAPI. **(c)** Immunoblot of synchronized HeLa cells expressing N100-E2F7-EGFP or N80-E2F8-mCherry. Cells were harvested following release from thymidine-nocodazole block. Cell cycle arrest/ progression was validated using anti-Securin antibodies (left) and DNA/ propidium iodide (PI) staining (right). Anti-Tubulin antibodies were used as a loading control. **(d)** Alignment of N100-E2F7 variants. Residue changes are highlighted (gray). **(e)** Time-dependent degradation of N100-E2F7 variants in S3 G1 extracts. Mean and SEM values are plotted (n = 3). The dynamics of N100-E2F7 and N80-E2F8 (a) are superimposed for comparison (dotted lines). **(f)** Time-dependent degradation of full-length E2F7 relative to E2F8 in S3 G1 extracts. **(g)** Time-dependent degradation of E2F7 relative to Securin in NDB mitotic extracts. (f,g) Quantifications of ^35^S signals (top) and representative source data (bottom) are shown (n = 3). Degradation assays were analyzed by SDS-PAGE and autoradiography.

Using a flow cytometry-based assay, we examined the temporal dynamics of EGFP-tagged N100-E2F7 and N80-E2F8 in vivo in asynchronous cells. Consistent with the degradation observed for N80-E2F8 in G1-phase extracts, the levels of N80-E2F8 dropped in G1-phase cells compared to other stages of the cell cycle. In contrast, N100-E2F7 levels, similar to the mCherry reporter, remained constant throughout the cell cycle **(Figures 2b)**. We next synchronized cells in mitosis and examined the dynamics of N100-E2F7 and N80-E2F8 after release from mitotic arrest into the cell cycle using western blot. This analysis confirmed a decline in N80-E2F8 levels, but not N100-E2F7, during the transition from prometaphase to G1 **(Figure 2c)**. Since the E2F8 KEN-box is located at the very N-terminus (residues 5-7), in contrast with the homologous E2F7 KEN-box (residues 31-33), we reasoned that the additional N-terminal residues in E2F7 might limit degradation in the artificial context of a short fragment **(Figure 2d)**. However, removal of residues 2-28 (E2F7Δ) did not restore degradation **(Figure 2e)**. KEN-box motifs are typified by a downstream proline residue [31] which is found in E2F8 (Pro 12) but is missing in E2F7 **(Figure 2d)**, suggesting a potential structural modification that might explain the difference in KEN-box functionalities observed in E2F7/8 fragments. However, insertion of an equivalent E2F7 proline (preceding residue Arg 38) did not restore degradation for N100-E2F7 (E2F7+P), nor for E2F7Δ (E2F7Δ+P) **(Figure 2e)**. We therefore conclude that the N-terminal fragment of E2F7, despite carrying a functional KEN motif, lacks critical elements required for APC/C-targeting. The molecular basis for this discrepancy remains unclear. Overall, the degradation of full-length E2F7 in G1 extracts was less effective compared to E2F8 **(Figure 2f)**, suggesting that the latter is more efficiently targeted by APC/C^Cdh1^.

We previously developed a mitotic extract system from HEK293T cells expressing a non-degradable mutant of Cyclin B1 (NDB) under a tetracycline (Tet)-inducible promoter [14, 32]. The persistent Cdk1 activity in this system prevents Cdh1 from binding to APC/C, thereby maintaining a consistent anaphase-like state with stable APC/C^Cdc20^ activity. Similar to E2F8 [14], E2F7 was stable in mitotic NDB extracts **(Figure 2g)**. Thus, the degradation patterns of E2F7 and E2F8 in mitotic- and G1 extracts are overall similar.

### Untimely degradation of non-phosphorylatable E2F8

Our previous findings demonstrated that phosphomimetic mutations at Thr20 and Thr44 of E2F8 (Thr to Asp) hampered degradation in HeLa G1 extracts. In contrast, the degradation of a non-phosphorylatable variant, where Thr20 and Thr44 are substituted with Ala, remained efficient [14]. These findings, suggest a role for the direct phosphorylation of Thr20 and Thr44 in regulating E2F8 half-life and highlight a broader connection between E2F8 (de)phosphorylation and degradation. To further investigate this, we examined the half-life of N80-E2F8 variants in NDB mitotic extracts. As previously reported, wild-type (WT) N80-E2F8 is stable in this extract system, which lacks APC/C^Cdh1^ activity **(Figure 3a** and Ref [14]). The phosphomimetic variant also remains stable in these extracts. In contrast, the Thr20/44-to-Ala double mutation destabilizes N80-E2F8 in a KEN-box-dependent manner **(Figure 3a)**, suggesting that this non-phosphorylatable variant can be targeted for degradation by APC/C^Cdc20^.

**Figure 3.**
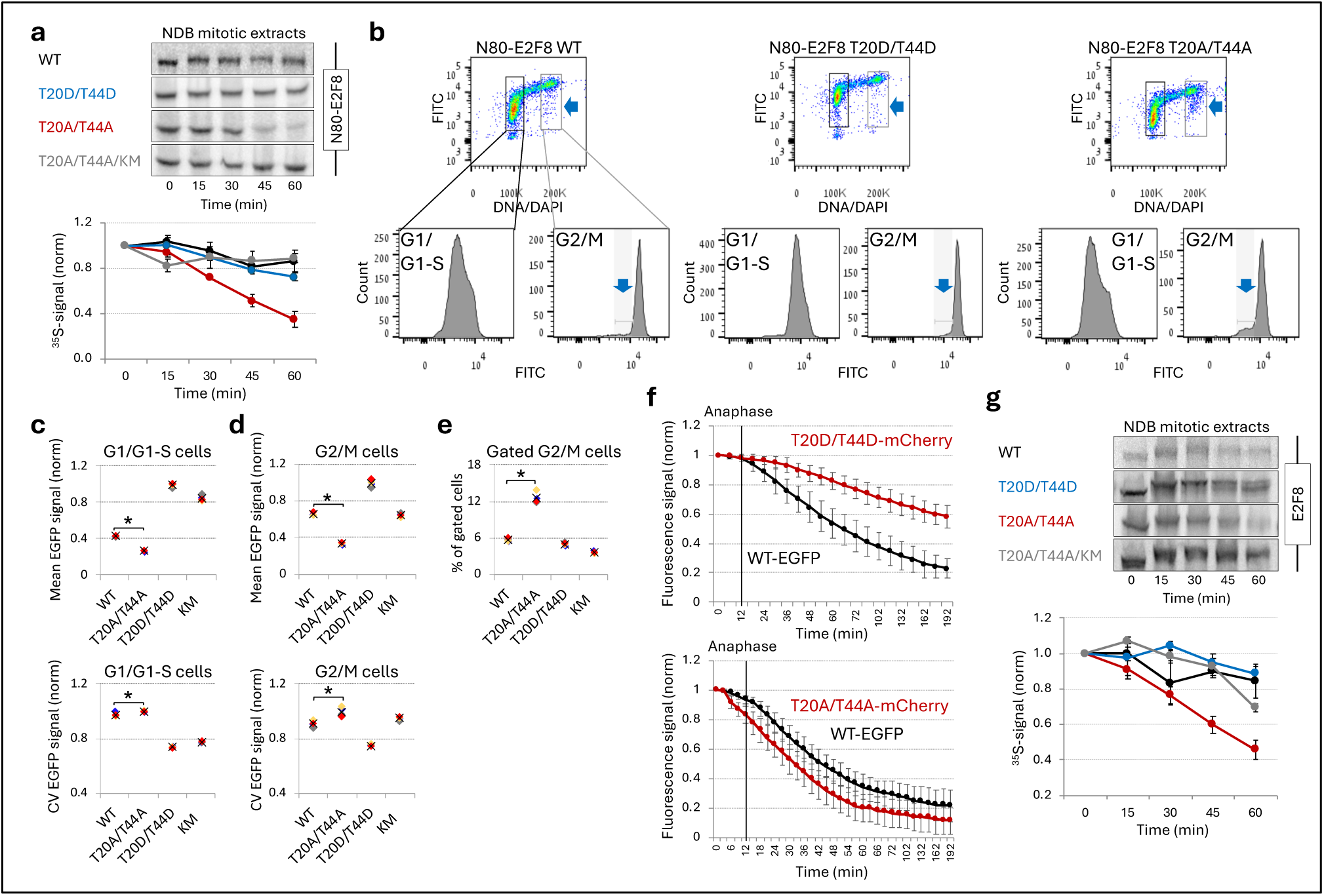
Non-phosphorylatable mutations induce untimely KEN-box-dependent degradation of E2F8. **(a)** Effect of phosphomimetic (Asp substitutions) and non-phosphorylatable (Ala substitutions) mutations at Thr20 and Thr44 on N80-E2F8 degradation in NDB mitotic extracts. The KEN-box mutant (KM) was generated by substituting the K-E-N sequence with A-A-N at positions 5-7 (single-letter amino acid codes). **(b)** Flow cytometry analysis of asynchronous HeLa cells expressing WT or mutant N80-E2F8-EGFP. Top: Bivariate plots showing DNA content (DAPI) versus EGFP fluorescence, with G1/G1-S phases and G2/M phases highlighted (black and gray frames, respectively). Bottom: Histograms showing EGFP signal distributions within these cell cycle phases. Blue arrows highlight subpopulations of G2/M cells with relatively low EGFP levels. **(c and d)** Quantification of EGFP signals in cells at G1/G1-S-(c) and G2/M-(d) phases. Mean (x) intensity and coefficient of variation (CV) are presented, comparing WT and mutant variants of N80-E2F8. Data are normalized; max mean value = 1. **(e)** Percentage of gated G2/M cells (from panel b) relative to the total G2/M cell population. (c-e) Experimental sets are color-coded. Mean = x; n = 4; *p < 0.05. **(f)** Time-lapse microscopy of HeLa cells co-expressing N80-E2F8-EGFP and either T20D/T44D or T20A/T44A variants tagged with mCherry. Mean fluorescence intensities and SDM are plotted (n = 20). Anaphase is indicated by a vertical line. **(g)** KEN-box-dependent degradation of the T20A/T44A E2F8 variant in NDB mitotic extracts. (a and g) Top: Representative data. Bottom: SEM values and normalized mean of ^35^S signal; n = 3. E2F8 variants are color-coded. Time-dependent degradation was analyzed by SDS-PAGE and autoradiography.

We next examined the steady-state distribution of EGFP-tagged N80-E2F8 variants in asynchronous cycling HeLa cells using flow cytometry, correlating EGFP signal with cell cycle phases as determined by DNA staining. The characteristic pattern of N80-E2F8 is shown in **Figure 3b** (top left plot), displaying the expected oscillations of an APC/C^Cdh1^ substrate: low in G1 phase, accumulating after APC/C is inactivated at the G1-S transition, and peaking in G2/M phases. A subset of G2/M cells exhibiting relatively lower EGFP signals (highlighted with blue arrows) is particularly informative for tracking APC/C^Cdc20^ substrates, such as Geminin, whose degradation begins in early mitosis, resulting in submaximal levels even in cells with 4N DNA that have not yet undergone cytokinesis [33].

This analysis was repeated for cells stably expressing phosphomimetic (T20D/T44D), non-phosphorylatable (T20A/T44A), and <u>K</u>EN-box <u>m</u>utant (KM) variants **(Figures 3b and S3a)**. The levels of phosphomimetic and KM mutants in G1/G1-S phases were significantly higher than those of WT N80-E2F8, as expected for more stable proteins **(Figure 3c, top)**. Correspondingly, the relative variability (CV) was low, indicating their limited dynamics **(Figure 3c, bottom)**. Interestingly, the T20D/T44D variant exhibited the highest EGFP level and the lowest CV, suggesting that the impact of these phosphomimetic mutations on E2F8 degradation is equal to or potentially surpasses that of a mutated KEN-box. Conversely, the non-phosphorylatable variant displayed significantly lower levels and higher CV in both G1/G1-S and G2/M phases, indicating heightened instability compared to WT N80-E2F8 **(Figures 3c and 3d)**. Furthermore, the percentage of G2/M cells with relatively low EGFP signals was more than two-fold higher for non-phosphorylatable N80-E2F8-EGFP compared to all other variants **(Figure 3e)**. Time-lapse analyses of HeLa cells co-expressing N80-E2F8-EGFP and either the phosphomimetic or non-phosphorylatable variants tagged with mCherry confirmed these findings; cells undergoing division maintained high levels of phosphomimetic N80-E2F8 even 3 hours after anaphase **(Figure 3f, top)**. More importantly, the decline of the non-phosphorylatable variant during mitotic exit preceded that of the WT protein and was more significant **(Figure 3f, bottom)**. Notably, degradation kinetics and HeLa time-lapse analyses of N80-E2F8 variants were independent of the fluorescent tag used **(Figures S3b and c)**, alleviating concerns of tag-specific effects.

We next examined the degradation of full-length E2F8 in NDB mitotic extracts. The non-phosphorylatable variant (T20A/T44A) was degraded in these extracts, while the WT and phosphomimetic variants remained stable **(Figure 3g)**. Moreover, the degradation was largely dependent on the ^5^KEN^7^ motif (KM, gray plot). Altogether, the data presented in Figure 3 demonstrate that non-phosphorylatable E2F8 is unstable during anaphase, can be targeted by APC/C^Cdc20^, and is rapidly degraded in cells exiting mitosis. Conversely, phosphomimetic E2F8 consistently remains stable during mitotic exit. These findings suggest that the phosphorylation status of Thr20 and Thr44 functions as a regulatory switch, determining the degradation of E2F8 throughout the cell cycle. At this juncture we note that attempts to visualize full-length E2F7/8 in cells, whether through knock-in tagging or stable ectopic expression, were unsuccessful, possibly due to cytotoxicity.

### E2F7 degradation in G1 extracts is modulated by N-terminal Cdk1 sites

Thr20 and Thr44 in E2F8 correspond to Thr45 and Thr68 in E2F7 **(Figure 4a)**. Indeed, alanine substitutions of Thr45 and Thr68 cooperatively reduced the Cdk1-dependent gel shift of N100-E2F7 following incubation in NDB mitotic extracts, potentially linking these residues to the overall regulation of E2F7 in cycling cells **(Figure 4b)**. In analogy to our above results for E2F8, we then examined the degradation of non-phosphorylatable (T45A/T68A) and phosphomimetic (T45D/T68D) E2F7 mutants in both mitotic and G1 extracts. Both WT and phosphomimetic variants remained stable in NDB mitotic extracts. Interestingly, and unlike E2F8, the Thr-to-Ala mutations did not induce degradation in such extracts **(Figure 4c)**. In contrast, the degradation of this variant in G1 extracts was accelerated **(Figure 4d)**, and in this case E2F8 produced comparable results **(Figure S4)**. Finally, phosphomimetic mutations hindered E2F7 degradation in G1 **(Figure 4d)**, consistent with previous findings for E2F8 [14]. Since Cdc20 activates APC/C in early anaphase, and Cdh1 activates APC/C during mitotic exit and G1 phase [11], these data imply that 1) phosphorylated E2F7/8 are resistant to APC/C^Cdc20^; 2) phosphorylation restricts E2F7/8 ubiquitination by APC/C^Cdh1^ in G1 phase; and 3) non-phosphorylatable forms of E2F7/8 are more efficiently targeted by APC/C^Cdh1^. Altogether, these results highlight a role for N-terminal Cdk1 sites in controlling the temporal dynamics of atypical E2Fs by modulating their APC/C-mediated ubiquitination.

**Figure 4.**
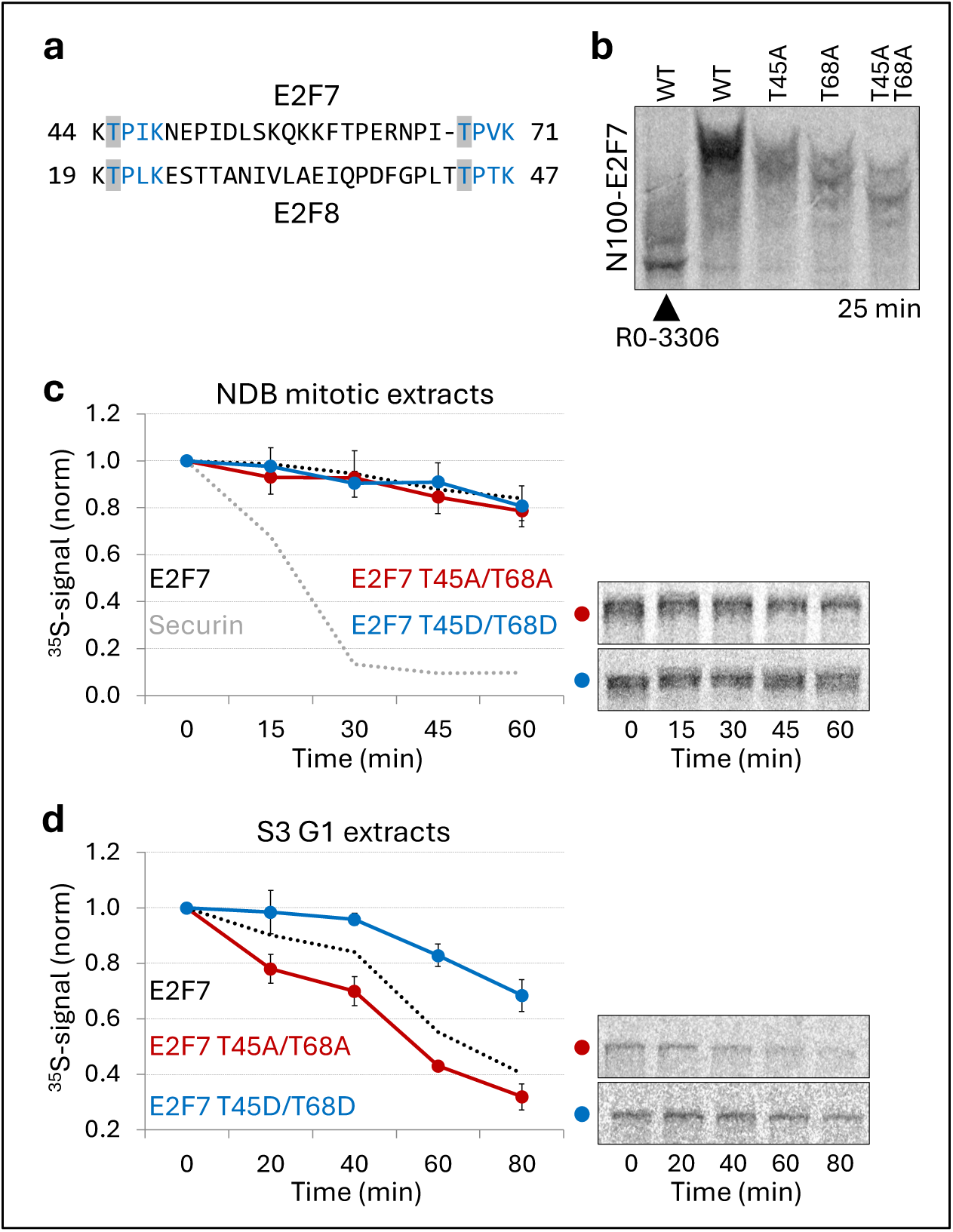
Phosphomimetic and non-phosphorylatable Cdk1 sites alter E2F7 degradation in G1 extracts. **(a)** BLAST alignment of E2F7/8 fragments, highlighting E2F7’s Cdk1 sites that correspond to Thr20 and Thr44 in E2F8. **(b)** Electromobility shift of N100-E2F7 variants in Phos-tag gel. IVT products were incubated in NDB mitotic extracts. RO-3306 was added to inhibit Cdk1-mediated shifts. **(c, d)** Time-dependent degradation of phosphomimetic and non-phosphorylatable variants of E2F7 in mitotic (c) and G1 (d) extracts. (c, d) Mean ^35^S signals SEM and are plotted (left); n = 3. Representative source data are shown (right). All assays were analyzed by SDS-PAGE and autoradiography.

### Cdk1 and PP2A control temporal (de)phosphorylation of E2F7 and E2F8

The N-terminal regions of E2F7/8 are enriched with Cdk1 consensus sites **(Figure 1a)**. To investigate the temporal (de)phosphorylation of these fragments throughout the cell cycle, we utilized cell-free systems recapitulating mitosis, mitotic exit, and various stages of S-phase [14]. Incubation of N100-E2F7 and N80-E2F8 in NBD mitotic extracts produced up to five distinct mobility-shifted forms, indicative of sequential phosphorylation events, all dependent on Cdk1 **(Figure 5a)**. Cdk1 inhibition (RO-3306) in NDB mitotic extracts triggers a switch from an active Cdk1-high mitotic state to a CdK1-low G1-like state [14, 32], leading to the gradual dephosphorylation of N100-E2F7 and N80-E2F8 **(Figure 5b)**. Phosphorylation was also evident in extracts derived from cells in mid-S phase **(Figure 5c)**, and was similarly inhibited by RO-3306, though not completely. Cdk2 is the predominant Cdk active during S phase, and RO-3306 likely inhibits both Cdk1 and Cdk2 [34]. These findings therefore suggest that E2F7/8 are targeted by Cdk2 in S phase, highlighting a role for Cdk2 in regulating atypical E2Fs through their N-terminal regions.

**Figure 5.**
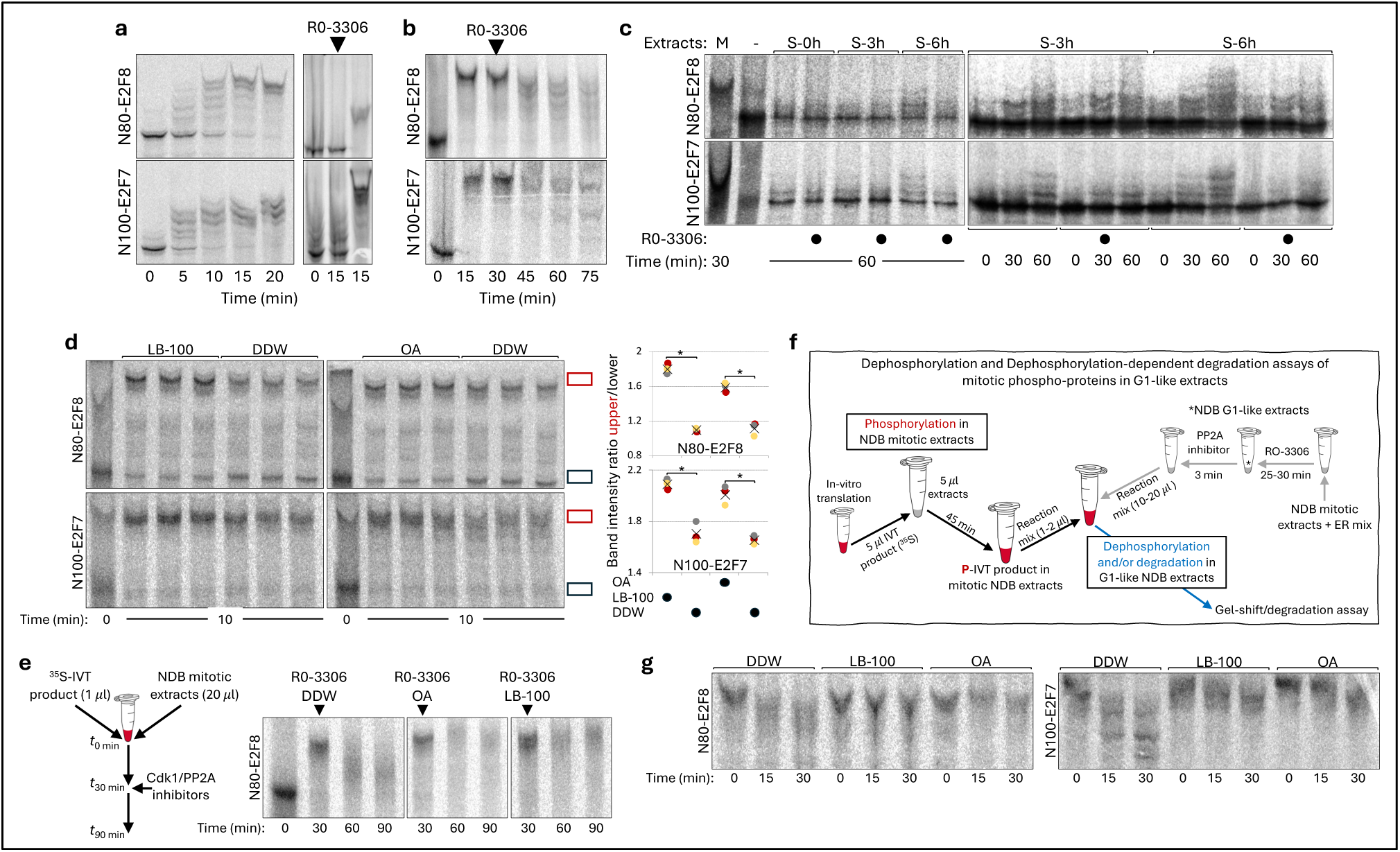
PP2A inhibitors block dephosphorylation of E2F7 and E2F8 during mitotic exit. **(a)** Left: Time-dependent phosphorylation of N80-E2F8 and N100-E2F7 during mitosis, indicated by electrophoretic mobility shifts following incubation in NDB mitotic extracts. Right: Equivalent assays where Cdk1 activity was inhibited by RO-3306. **(b)** Dephosphorylation of N80-E2F8 and N100-E2F7 during mitotic exit. Mitotic extracts, pre-mixed with IVT products for 30 minutes, were induced to exit mitosis by RO-3306. **(c)** Gel-shift assays of N80-E2F8 and N100-E2F7 after incubation in NDB mitotic extracts (M) and S-phase extracts prepared from synchronous HeLa S3 cells harvested at 0, 3, and 6 hours post-release from a double thymidine block (S-0/3/6 h). Both end-point (left) and time-dependent (right) assays are shown, with Cdk1/2 activity blocked by RO-3306. **(d)** Gel-shift assay showing enhanced phosphorylation of E2F7/8 in mitotic extracts treated with LB-100 or Okadaic acid (OA). The ratio between fully phosphorylated and unphosphorylated forms of N80-E2F8 and N100-E2F7 (left; marked by red and black rectangles, respectively) was plotted (right). Charts present mean (x) and individual data points (n = 3). *p < 0.05. **(e)** (De)phosphorylation of N80-E2F8 in mitotic extracts co-treated with Cdk1 and PP2A inhibitors (right). Left: Experimental strategy. **(f)** Schematics of a method to detect both dephosphorylation and dephosphorylation-dependent degradation of IVT products in cell-free systems. **(g)** Inhibition of E2F7/8 dephosphorylation in NDB G1-like extracts by PP2A inhibitors. Target proteins were pre-phosphorylated in NDB mitotic extracts and then shifted to G1-like extracts containing PP2A inhibitors (detailed in panel f). Overall, the assays were performed using ^35^S-labeled IVT products. Reaction mixtures were resolved by Phos-tag SDS-PAGE and analyzed by autoradiography. Drug concentrations: RO-3306, 30 µM; OA, 2.5 µM; LB-100, 500 µM.

Phosphatases countering the action of Cdk1/2 are likely to play a significant role in the overall regulation of the E2F1-E2F7/8 network. The holoenzymes PP1 and PP2A are the major CdK1-counteracting phosphatases during mitotic exit, making them candidate regulators of E2F7/8. To test this hypothesis, N100-E2F7 and N80-E2F8 underwent phosphorylation in NDB mitotic extracts in the presence of PP2A inhibitor LB-100 [35, 36] and PP1/PP2A inhibitor okadaic acid (OA). This manipulation enhanced the mitotic phosphorylation of E2F7/8 **(Figure 5d)**. In turn, the dephosphorylation of N80-E2F8 was significantly blocked in extracts co-treated with both Cdk1 and PP1/PP2A inhibitors, further solidifying a potential link between E2F7/8 and PP1/PP2A **(Figure 5e)**. However, the inhibition of pleiotropic phosphatases such as PP2A may cause non-specific distortion of mitotic exit. To address this concern, we developed a direct dephosphorylation assay in which NDB mitotic extracts containing pre-phosphorylated targets are mixed in a ∼1:10 v/v ratio with NDB extracts in a G1-like state induced by RO-3306 (**Figure 5f**). This allows for direct analysis of dephosphorylation as well as dephosphorylation-dependent degradation of exogenously added proteins in a human cell-free system. The addition of OA and LB-100 to this system significantly blocked dephosphorylation of both N80-E2F8 and N100-E2F7 **(Figure 5g)**. These results demonstrate that the dephosphorylation status of E2F7/8 in G1 phase is dependent on PP2A and collectively suggest that the temporal control of E2F7/8 phosphorylation is balanced by the opposing activities of Cdk1, Cdk2, and PP2A.

### APC/C-mediated degradation of E2F7 and E2F8 relies on a dephosphorylation switch

We next determined if the regulation of E2F7/8 by the APC/C is directly impacted by (de)phosphorylation. To address this question, we tested the half-life of phosphorylated (P) vs. unphosphorylated (UP) E2F7/8 in G1 extracts supplemented with 250 µM LB-100. At this concentration, dephosphorylation of the mitotic form of N80-E2F8 is severely blocked **(Figure 6a)**. Although APC/C-mediated degradation is also affected, it remains efficient, allowing us to measure relative half-lives of APC/C targets that cannot undergo PP2A-mediated dephosphorylation. Degradation assays for phosphorylated targets required in vitro translated proteins (IVT) that were pre-mixed with NDB mitotic extracts and performed in G1-like extracts (RO-3306-activated) derived from NDB cells **(Figures 5f and 6b)**. The degradation of both the phosphorylated N80 fragment and full-length E2F8 in NDB G1 extracts was significantly impaired compared to the unmodified proteins **(Figure 6c)**. Moreover, the degradation of non-phosphorylatable variants of N80-E2F8 and E2F8 (T20A/T44A) was only moderately affected by mitotic phosphorylation **(Figure 6d)**.

**Figure 6.**
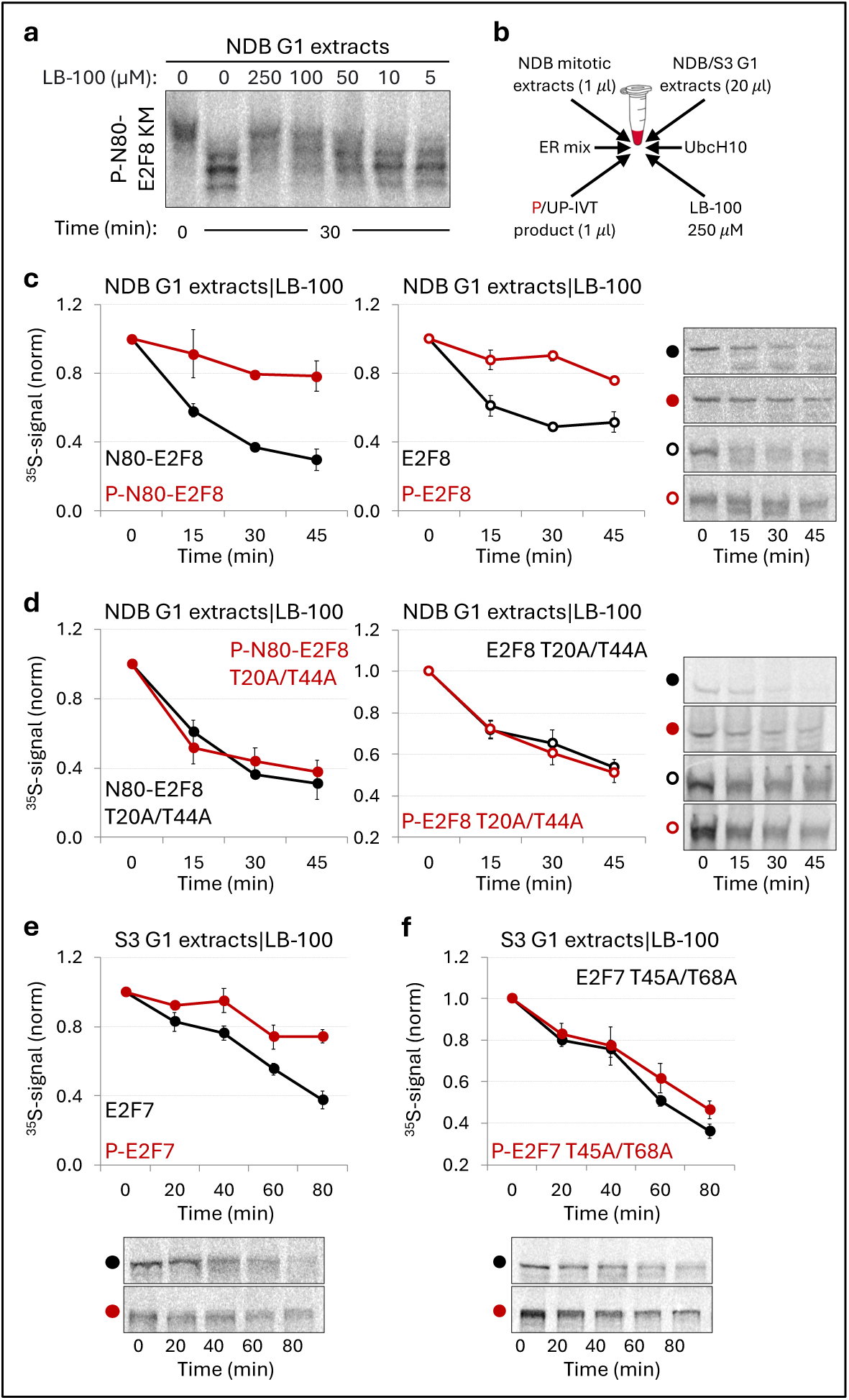
G1 degradation of E2F7 and E2F8 is controlled by PP2A-mediated dephosphorylation. **(a)** Dose-dependent inhibition of PP2A by LB-100, as assessed by Phos-tag gel shifts of a phosphorylated (P)-variant (KEN-box mutant) of N80-E2F8. The IVT product was pre-phosphorylated in NDB mitotic extracts, then a portion of the mixture was incubated in NDB G1-like extracts with increasing doses of LB-100 for 30 minutes. **(b)** Schematic for detecting dephosphorylation-dependent degradation of mitotic (<u>u</u>n)<u>p</u>hosphorylated (UP/P) targets. **(c, d)** Time-dependent degradation of P/UP N80-E2F8 (left) and full-length E2F8 (right) in NDB G1 extracts, with assays for both WT (c) and non-phosphorylatable (d) variants. **(e, f)** Time-dependent degradation of P/UP E2F7 in S3 G1 extracts, comparing WT (e) and non-phosphorylatable (f) variants. All assays were analyzed by SDS-PAGE and autoradiography, with mean ^35^S signal and SEM (n = 3) plotted alongside representative source data.

The reaction conditions used to analyze E2F8 degradation **(Figure 6d)** did not support meaningful degradation of E2F7 **(Figure S5)**. However, G1 extracts from HeLa S3 cells revealed reduced degradation rates for mitotic-phosphorylated E2F7 as well **(Figure 6e)**. Consistent with E2F8, the degradation of the Thr45/68-to-Ala E2F7 mutant was nearly unaffected by mitotic phosphorylation **(Figure 6f)**. Together, these results underscore PP2A-mediated dephosphorylation as a critical molecular switch in controlling the degradation of atypical E2Fs. Furthermore, they suggest that the phosphorylation state of key Cdk1 sites in the unstructured N-terminal region serves as a molecular integrator, regulating the timely degradation of both transcription factors in cycling mammalian cells.

## Discussion

In this work we demonstrate a dephosphorylation switch that couples the proteasomal degradation of atypical E2Fs to mitotic exit and the G1 phase **(illustrated in Figure 7a)**. Key Thr residues in the N-terminal regions of E2F7/8 are phosphorylated by Cdk1 and likely Cdk2. While this phosphorylation may facilitate ubiquitination via the SCF ligase, which is known to target phospho-degrons, it prevents premature degradation, at least for E2F8 **(Figure 3)**. Upon mitotic exit, PP2A dephosphorylates E2F7/8, rendering them susceptible to degradation via APC/C^Cdh1^. Concomitantly, PP2A facilitates the transition from APC/C^Cdc20^ to APC/C^Cdh1^ [18]. Thus, the counteractivity between Cdk1 and PP2A serves as a dual safety mechanism, ensuring the timely degradation of atypical E2Fs.

**Figure 7.**
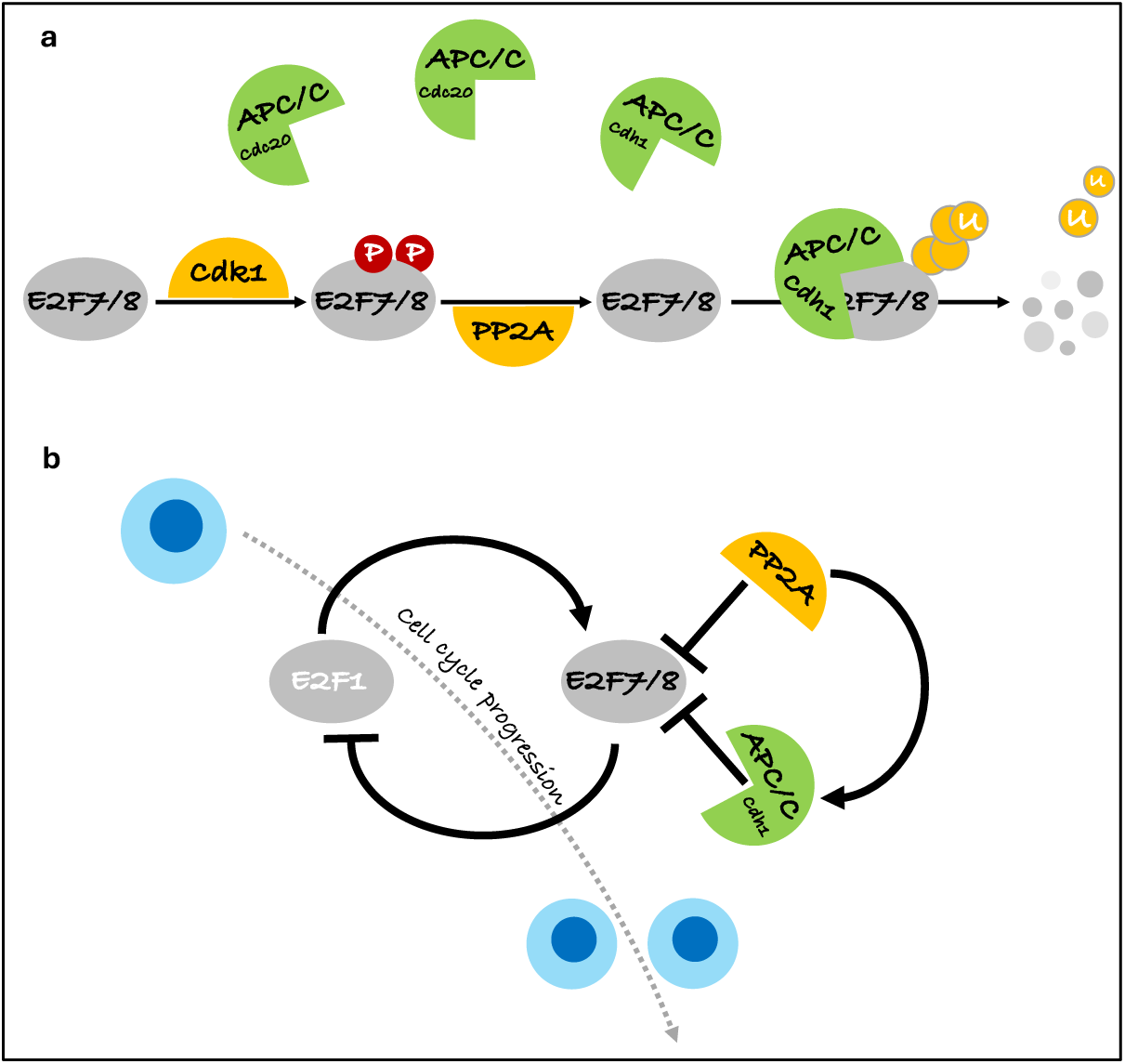
Cdk1-PP2A counteractivity establishes molecular switches that modulate atypical E2F degradation. The interplay between Cdk1 and PP2A activities ties E2F7/8 degradation to the cell-cycle clock. During mitosis, Cdk1 phosphorylates the disordered N-terminal regions of E2F7/8, preventing premature degradation. As cells transition to anaphase, the decrease in Cdk1 activity allows PP2A to dephosphorylate E2F7/8 at key Thr residues, thereby promoting degradation by APC/C^Cdh1^. Additionally, PP2A targets Cdh1, enhancing its interaction with APC/C. This critical step facilitates the shift from APC/C^Cdc20^ to APC/C^Cdh1^ during mitotic exit **(a)**. The regulatory axis described here, involving feedback and crosstalk between Cdk1, PP2A, and APC/C, integrates into the broader regulation of the mammalian cell cycle through the E2F1-E2F7/8 transcriptional network (b)

We have previously noted that the degradation of phosphomimetic E2F8 variants in G1 extracts is relatively inefficient [14], and here, a similar phenotype is observed for a homologous E2F7 mutation **(Figure 4)**. Although phosphomimetic mutations are useful for studying protein regulation by phosphorylation, there is a concern that they introduce structural changes beyond merely mimicking the phosphate group. In the current work, a new cell-free assay analyzing dephosphorylation-dependent degradation allowed us to measure the half-life of Cdk1-phosphorylated targets in a G1 environment with inhibited PP2A activity, without interfering mitotic exit **(Figure 5f)**. Our results indicate that APC/C^Cdh1^-mediated degradation of both E2F7/8 is regulated by PP2A counteracting Cdk1-dependent phosphorylation **(Figure 6)**. Given that E2F7/8 are targeted by S-phase kinases, likely Cdk2 **(Figure 5c)**, it is reasonable to speculate that the opposing actions of Cdk1/2 and PP2A serve to confine E2F7/8 degradation. The limited and accelerated degradation rates observed for phospho-mimetic and non-phosphorylatable variants of E2F7/8 in G1, respectively, align well with this model **(Figures 3, 4 and S4)**. Thus, (de)phosphorylation appears to directly modulate both the E3 ligase (APC/C) and its substrates, ensuring that E2F1-driven transcription is neither prematurely restrained nor overly prolonged once cells have entered S-phase. The catalytic subunit of PP2A is known to interact with numerous regulatory subunits, particularly the B55 and B56 types during mitotic exit [37]. While the exact PP2A holoenzyme responsible for targeting E2F7/8 has not been explored here, this raises compelling questions for further investigation.

Neither E2F8 nor E2F7 is degraded in NDB mitotic extracts, suggesting that they are not targeted for destruction by APC/C^Cdc20^. APC/C substrates typically contain multiple degradation motifs [38]. However, ^5^KEN^7^ is the only known APC/C recognition element in N80-E2F8. KEN-box motifs are recognition elements predominantly associated with APC/C^Cdh1^, unlike the canonical destruction-box (D-box) motif, which is recognized by both Cdc20 and Cdh1 [38–40]. Surprisingly, non-phosphorylatable variants of E2F8 exhibited significant instability in NDB mitotic extracts, which was KEN-box-dependent **(Figure 3)**. Evidence for KEN-box substrates of APC/C^Cdc20^ is scarce but has been reported, as seen with the centrosomal kinase Nek2A [41]. Overall, our findings suggest that the Cdc20/Cdh1 specificity of E2F8, and consequently its degradation timing, is determined by both integrated degradation signals and conditional phosphorylation status.

Sequence and NMR-based structural analyses suggest that the N-terminal regions of E2F7/8 are highly flexible and adopt a disordered and unfolded conformation **(Figure 1)**. Such flexibility renders the polypeptide sequence as ‘clay in the potter’s hands’ since disordered regions are more susceptible to conformational changes induced by post-translational modifications [42, 43]. This model is consistent with the role of unfolded N-terminal regions as phospho-regulatory degron domains, akin to those found in other APC/C substrates [28, 29, 44–46].

The transcriptional activity of E2F7/8 relies on homo- and heterodimerization; they share overlapping and redundant roles, particularly in repressing E2F1 transcription [7]; and are co-regulated by similar mechanisms, leading to comparable spatiotemporal dynamics and fates [11]. Our results extend these parallels by demonstrating dephosphorylation-mediated degradation for both E2F7 and E2F8 **(Figure 6)**, along with low structural propensities of their N-terminal domains **(Figure 1)**. However, we also highlight notable differences between these proteins. First, E2F7 possesses an unstructured N-terminal tail with an unknown function **(Figure 1)**. Second, the N-terminal segment of E2F7 is not targeted by APC/C, despite carrying a degradation signal **(Figure 2a-e)**. Third, E2F7 degradation in G1 extracts appears less efficient compared to E2F8, possibly due to weaker APC/C^Cdh1^ substrate processivity and/or binding **(Figure 2f)**. Lastly, only the non-phosphorylatable variant of E2F8 exhibits instability in mitotic extracts **(Figures 3 and 4)**. The biological relevance of these distinctions remains to be elucidated.

Our findings are situated within three core negative feedback mechanisms that regulate the cell cycle **(illustrated in Figure 7b)**. E2F7/8 cooperate to downregulate E2F1, balancing E2F1-mediated transcription following S-phase entry. At the post-translational level, Cdk1 and PP2A engage in reciprocal negative regulation, orchestrating mitotic entry and transitions from mitosis to the G1 phase. Additionally, APC/C^Cdh1^ targets E2F7/8 for degradation, allowing E2F1 to establish the molecular framework necessary for the S- and M-phases. This includes Emi1 [47] and Cyclin A [48], which coordinate the G1-S transition by inhibiting and dismantling APC/C^Cdh1^ itself [49, 50]. Collectively, our results demonstrate a nuanced interplay between phosphorylation and degradation, highlighting a layer of regulation that fine-tunes the temporal activity of the E2F1-E2F7/8 circuitry in governing the transcriptional program of the mammalian cell cycle.

## Materials and Methods

### Plasmids

Table S1 provides a list of all plasmids used in this study. Cloning strategies are detailed for newly generated plasmids, while previously published plasmids are considered lab stock. Cloning and mutagenesis procedures were validated by Sanger sequencing.

### Cell culture

The cell lines used in this study are HeLa, HeLa S3 (S3), and HEK293T, originally obtained from the American Type Culture Collection (ATCC). Cells were cultured in DMEM supplemented with 9% fetal bovine serum, 2 mM L-glutamine, and 1% penicillin-streptomycin solution (Biological Industries; #03-020-1B, #03-031-1B). Cells were maintained at 37°C in a humidified atmosphere containing 5% CO_2_. HeLa cell lines generated by stable viral infection include: 1) N80-E2F8-EGFP; 2) N80-E2F8-T20A/T44A-EGFP; 3) N80-E2F8-T20D/T44D-EGFP; 4) N80-E2F8-KM-EGFP; 5) N80-E2F8-T20A/T44A-EGFP and N80-E2F8-mCherry; 6) N80-E2F8-T20D/T44D-EGFP and N80-E2F8-mCherry; 7) N80-E2F8-T20A/T44A-mCherry and N80-E2F8-EGFP; 8) N80-E2F8-T20D/T44D-mCherry and N80-E2F8-EGFP; 9) N100-E2F7-EGFP and mCherry. Lentiviral particles were produced in HEK293T cells co-transfected (calcium phosphate) with PTZV vectors, into which the relevant expression sequences were cloned, alongside additional plasmids encoding, Rev, VSV-G, and Gag-Pol. Stably infected cell lines were derived from single-cell colonies and selected with 2 µg/ml puromycin and/or by fluorescence signals. The generation of NDB cells used for cell-free systems is described in Ref [14, 32]. NDB cells were maintained in a selection medium containing 200 µg/ml Zeocin (Invitrogen; # R25001) and 5 µg/ml Blasticidin (Life Technologies; #A11139-03).

### Reagents

The following reagents were used: 1) Thymidine, 2 mM (Sigma-Aldrich, #T9250); 2) Nocodazole, 50 ng/ml (Biotrend, #A8487); 3) Tetracycline, 1 µg/ml (Sigma-Aldrich, #87128); 4) PP1/PP2A inhibitor Okadaic Acid, 2.5 µM (Enzo Life Sciences, #ALX-350-003-C025); 5) PP2A inhibitor LB-100, 250 µM default concentration (Abcam, #ab285402); 6) Cdk1 inhibitor RO-3306, 30 µM (Focus Biomolecules, #10-4126); and 7) MG-132 proteasome inhibitor, 30 µM (Tocris, #1748).

### Cell synchronization

#### HeLa S3 cells for G1 extracts

HeLa S3 cells were cultured in a 1L spinner flask at 85 rpm for 2-3 days until reaching a density of 4 × 10^5^ cells/ml. Thymidine was added for 22 hours to synchronize the cells in S phase. Afterward, the cells were washed and re-cultured in fresh, pre-warmed media for 3 hours. Subsequently, nocodazole was added for 12 hours to induce mitotic arrest. Cells were washed, re-cultured in pre-warmed fresh media, and harvested after 3.5 hours.

#### HeLa cells for immunoblot assays

HeLa cells were cultured in 15 cm diameter dishes and were at 70% confluency when Thymidine was added to initiate a Thymidine-nocodazole block (see previous section). Cells were harvested via mitotic shake-off and either collected immediately (t_0_) or recultured for 4 to 7 hours before harvesting.

#### HeLa S3 cells for S-phase extracts

S-phase extracts were generated from HeLa S3 cells harvested at 0-, 3-, and 6-hours post-release from a double-thymidine block. The synchronization protocol is detailed in Ref [14].

#### NDB HEK233 cells for mitotic extracts

NDB HEK293 cells were cultured to 80% confluency and treated with Tetracycline for 20-22 hours before harvesting. Cell synchronization was validated using Propidium Iodide (PI; Sigma-Aldrich, #P4170).

### Expression and purification of N100-E2F7 and N80-E2F8

N100-E2F7 and N80-E2F8 were expressed as maltose-binding protein (MBP) fusion constructs, including a His_6_-tag and a tobacco etch virus (TEV) protease cleavage sequence (MBP-His_6_-TEV-E2F7/8), in BL21 *E. coli* cells. Purification followed a standardized protocol. Cells were grown in M9 minimal medium containing ¹⁵NH₄Cl (1 g/L), 2.5 g/L ¹³C₆-D-glucose (Cambridge isotope laboratories, #NLM-467-PK and CLM-1396-PK), and 0.25 g/L of the ¹³C,¹⁵N Isogro supplement (Sigma-Aldrich, #606839). At an OD₆₀₀ of approximately 0.8, cultures were induced with 1 mM IPTG and expressed overnight at 27 °C. After expression, cells were pelleted by centrifugation at 8000 rpm (Beckman JA-14 rotor) for 10 minutes at 4 °C, followed by lysis using a C5 homogenizer (Avestin) in lysis buffer (500 mM NaCl, 10 mM Tris, pH 7.5, 10 mM imidazole, 1 mM DTT) supplemented with one tablet of EDTA-free protease inhibitor cocktail and 5 mM benzamidine. Following homogenization, additional benzamidine (final concentration 10 mM) and phenylmethylsulfonyl fluoride (PMSF, final concentration 1 mM) were added. The lysate was clarified by centrifugation at 18,000 rpm for 40 minutes at 4 °C. The supernatant was loaded onto a Ni²⁺-affinity column, and the protein was eluted with 50 mM imidazole. To the eluted fraction, one tablet of EDTA-free protease inhibitor cocktail and 5 mM benzamidine were added. MBP cleavage was performed using 1:20 w/w TEV protease for 3 hours at 30 °C. The mixture was then reloaded onto the Ni²⁺-affinity column, and the flow-through fraction containing the target polypeptide was collected. This eluted fraction was acidified with 1 M Tris-formate buffer to pH 3.5 to prevent aggregation, desalted by reverse phase (RP) chromatography, and lyophilized. For NMR measurements, samples were reconstituted in 20 mM HEPES, pH 7.5, 5 mM KCl, 2 mM MgCl₂, 1 mM DTT, one tablet of EDTA-free protease inhibitor, and 5% ²H₂O. Measurements were conducted at 16.4 T and 278 K.

### NMR spectroscopy for N100-E2F7 and N80-E2F8

All 2D and 3D NMR measurements were conducted on a DRX700 Bruker spectrometer using a cryogenic triple-resonance TCI probehead equipped with z-axis pulsed field gradients. Assignment of resonances was achieved by acquiring standard backbone triple-resonance experiments, including HNCO, HNCA, HNCOCAB, and HNCACB [51], and was aided by a ¹³C’-detected strategy (the 2D ¹³C’-¹⁵N correlation spectrum as readout) [52]. This strategy included 3D experiments CANCO, CBCACON, and CBCANCO, which are suitable for disordered proteins with poor spectral dispersion. Backbone resonances of both proteins were used as input for the delta2D program [26] to predict secondary structure content. R₁ρ relaxation rates for both proteins were measured as the decay in a 2D ¹H,¹⁵N-HSQC spectrum in an interleaved fashion, with relaxation delays of 0.016, 0.112, 0.208, 0.304, and 0.400 seconds, and were analyzed using the DYNAMICS program suite.

Sequence alignment was conducted using BLASTp. Prediction of disordered regions was performed with the IUPRED3 program [25], employing a cutoff of 0.5 for the disorder fraction.

### Cell Imaging and Flow Cytometry

#### Live-cell microscopy and image analysis

Live cells were imaged using the IncuCyte® live-imaging system (Sartorius). ImageJ (National Institutes of Health) software was employed for image processing and analysis, including manual cell segmentation and tracking.

#### Flow Cytometry

The LSRFortessa™ cell analyzer and FACSAria™ III cell sorter (BD Biosciences) were utilized for analysis and single-cell sorting, respectively. Analysis of DNA and EGFP/mCherry signals was performed on cells fixed with 4% paraformaldehyde (PFA). DNA was probed with DAPI stain (5 µg/ml; RayBiotech, #331-30016) at room temperature for 10 minutes. Data processing and analysis were conducted using FlowJo™ software (BD Biosciences).

### Immunoblotting

Cells were washed twice in PBS and lysed for 30 minutes on ice. The lysis buffer (50 mM Tris, pH 7.6, 150 mM NaCl, 5 mM EDTA, pH 8.0, 0.5% NP-40) was supplemented with: 1) protease inhibitor cocktail (Roche; #11836170001); 2) 1 mM PMSF; 3) phosphatase inhibitor cocktail (Sigma-Aldrich; #P5726, #P0044); 4) 10 mM NaF; 5) 20 mM β-glycerophosphate; and 6) 1 mM Na₃VO₄. The lysed cells were centrifuged at 14,000 × g for 40 minutes. Protein lysate concentration was measured using Bradford reagent (Bio-Rad; #500-0006) and Epoch spectrophotometer. Protein samples were mixed with Laemmli buffer, denatured (7 minutes, 95 °C), and resolved by sodium dodecyl sulfate polyacrylamide gel electrophoresis (SDS-PAGE). Proteins were electro-transferred onto a nitrocellulose membrane Nupore; #SCN02279.4). Protein loading and transfer quality were verified using Ponceau S staining solution (Sigma-Aldrich; #P7170). Membranes were blocked with 5% skimmed milk in TBST for 1 hour and then incubated overnight at 4 °C with the following primary antibodies: 1) anti-GFP (Santa Cruz Biotechnology; #SC9996); 2) anti-mCherry (Abcam, #ab125096); 3) anti-Securin (Cell Signaling Technology; #13445); and 4) anti-Tubulin (Sigma-Aldrich; #T9026). Antibodies were diluted 1:1000 in antibody solution (4% BSA and 0.05% sodium azide in TBS). Secondary antibodies conjugated with horseradish peroxidase were purchased from Jackson ImmunoResearch (#115-035-003, #111-005-003). The ECL reaction was performed using EZ-ECL (Thermo Scientific; #34580) and analyzed with the Amersham Imager 680 (GE Healthcare).

### Preparation of cell extracts

#### HeLa S3 extracts

A 400-500 mL culture of synchronous HeLa S3 cells was washed twice with cold PBS, lysed in swelling buffer (20 mM HEPES, pH 7.5, 2 mM MgCl₂, 5 mM KCl, 1 mM dithiothreitol [DTT], and protease inhibitor cocktail [Roche; #11836170001]), and incubated on ice for 30 minutes. The lysed cells were homogenized by freeze– thawing in liquid nitrogen and passed through a 21-G needle 10 times for mechanical shearing in a native environment. Extracts were cleared by two rounds of centrifugation (14,000 RPM, 10 and 40 minutes) and supplemented with an energy regenerating (ER) mix (1 mM ATP, 0.1 mM ethylene glycol-bis (β-aminoethyl ether)-N,N,N′,N′-tetraacetic acid [EGTA], 1 mM MgCl₂, 7.5 mM creatine phosphate, and 50 µg/mL creatine phosphokinase). Cell extracts were snap-frozen in 45 µL aliquots in liquid nitrogen and stored at −80 °C, with each aliquot thawed only once. Protein concentration in the extracts was 20 ± 3 mg/mL.

#### NDB mitotic extracts

NDB cells at 80% apparent confluency were treated with Tetracycline for 20-22 hours. At this stage, nearly all cells are in an anaphase-like state, either floating or loosely attached to the surface, allowing for direct harvesting for the preparation of mitotic extracts (see preceding paragraph). NDB mitotic extracts are typically generated from 20 plates of 15 cm diameter. Protein concentration in the extracts was 25 ± 3 mg/mL.

#### NDB G1-like extracts

NDB mitotic extracts can be induced to exit mitosis into G1 phase by adding 25-30 µM RO-3306 along with ER mix. After 25-30 minutes at 28-30 °C, the extracts are prepared for assessing PP2A-mediated dephosphorylation and APC/C^Cdh1^-mediated degradation of target IVT proteins. For more information, see Ref [14, 32].

### Protein degradation assay in cell extracts

Target proteins were in vitro transcribed and translated using the SP6 TNT Quick coupled reticulocyte system (Promega; #L2080), supplemented with a ³⁵S-methionine/cysteine cocktail (PerkinElmer; #NEG772002MC). The standard degradation assay consists 20 µL of cell extract, 1 µL of IVT product, 1 µL of ER mix, and 1 µL of recombinant His-tagged UbcH10 or UbcH10DN (10 µM). When applicable, small-molecule inhibitors were also added. The final reaction volume was typically 23-24 µL. Time-dependent degradation assays were performed in 0.2 mL PCR tubes at 28-30 °C, with samples collected directly into 10 µL of Laemmli buffer, followed by snap-freezing in liquid nitrogen and denaturation at 95 °C for 7 minutes. Sample volumes of 3-5 µL were sufficient to generate a readable signal. Proteins were resolved by SDS-PAGE. Gels were soaked in a destaining solution (10% methanol, 7.5% acetic acid) for 20 minutes, vacuum-dried (Gel Dryer Model 583, Bio-Rad), and exposed to a phosphor screen overnight or longer. IVT proteins were visualized by autoradiography using a Typhoon FLA 9500 phosphorimager (GE Healthcare Life Sciences). Signal intensity was quantified using ImageJ software.

### Phosphorylation assays

NDB mitotic extracts maintained high phosphorylation activity, even when diluted 10-fold, allowing efficient phosphorylation at a high substrate/extract volume ratio of 1:1. In a typical assay, 1 µL of IVT substrate (³⁵S-labeled) is mixed with 10-20 µL of undiluted NDB mitotic extracts and 1 µL of ER mix. The reaction temperature ranges between 23 to 28 °C. Reaction samples are aliquoted for each time point and processed as described in the preceding paragraph. Proteins are resolved on freshly made 8% acrylamide gels containing 20 µM PhosTag reagent (APExBIO; #F4002), which enhances electromobility shifts in SDS-PAGE. Gels are fixed, vacuum-dried, and analyzed by autoradiography.

### Dephosphorylation/dephosphorylation-dependent-degradation assays

These assays are performed on mitotically phosphorylated IVT targets (³⁵S-labeled). For pre-phosphorylation, IVT proteins are incubated with NDB mitotic extracts (v/v = 1) and ER mix for 45 minutes at 28 °C. Phosphorylation is validated by gel-shift assays (PhosTag). Importantly, the high substrate concentration makes IVT targets detectable even when diluted 50-fold, circumventing the need to purify the phosphorylated target protein from the mitotic extracts for further assays. Dephosphorylation is assayed in HeLa S3/NDB G1 extracts. To this end, 1-2 µL of the mitotic reaction solution is mixed with 10-20 µL of G1 extracts. The 5-10% mitotic activity does not jeopardize the PP2A and APC/C^Cdh1^ activities. Time-dependent dephosphorylation is analyzed by PhosTag SDS-PAGE. When analyzing dephosphorylation-dependent degradation, 250 µM LB-100 is added to block PP2A-mediated dephosphorylation of the tested proteins during incubation in G1 extracts. Control assays testing unphosphorylated targets are also supplemented with LB-100 and mitotic NDB extracts to equalize reaction conditions. Degradation in LB-100-treated extracts is attenuated yet remains potent and informative. Time-dependent degradation is analyzed by SDS-PAGE and autoradiography.

### Statistical analysis

A Student’s t-test (two-tailed) was used to determine statistical significance. Degradation assays and time-lapse microscopy data are presented as mean ± SEM/SDM. The number of replicates (n) and p values are provided in the figure legends, with *p < 0.05 indicating significance. Microsoft Excel was used for creating plots and performing statistical analysis.

## Acknowledgments

This work was supported by the Israel Science Foundation; Grant Nos. 2038/19, 1356/24, and 1362/24 (Tzur lab), and 964/19 (Chill lab). We also acknowledge support for the Emanuele lab from the National Institutes of Health; Grant no. R35GM153250.

