## Supplementary Figures and Figure legends for "Cdk1 and PP2A constitute a molecular switch controlling orderly degradation of atypical E2Fs": Supp.pdf

**Title:**

**Notes:**

[*] Corresponding author

Electronic address:

**Keywords:**

Cell cycle, Mitotic exit, E2F, Ubiquitination, APC/C substrate, Cell-free system, Protein degradation

**Abstract**

Dynamic oscillations in the phosphorylation and ubiquitination of key proliferative regulators are defining features of the eukaryotic cell cycle. Resetting the cell cycle at the mitosis-to-G1 transition requires activation of the E3 ubiquitin ligase Anaphase-Promoting Complex/Cyclosome (APC/C), which ensures cell cycle irreversibility by targeting dozens of substrates for degradation, safeguarding genome integrity. However, the overall coupling of substrate phosphorylation with target recognition and degradation by the APC/C remains relatively unexplored. As a paradigm for further defining these rules, we focused on E2F7 and E2F8 – atypical E2F-family proteins that coordinate cell cycle gene expression by restraining the pro-proliferative transcriptional activity of E2F1. Leveraging complementary cell and cell-free systems, we demonstrate that flexible domains in the amino-termini of E2F7 and E2F8 contain APC/C recognition motifs adjacent to critical Thr residues, whose phosphorylation by Cdk1 is rate limiting for degradation. The removal of this phosphorylation by PP2A phosphatase serves as a molecular switch, coupling the degradation of E2F7 and E2F8 to the G1 phase, coinciding with the rise of E2F1. Collectively, these findings highlight a critical role for Cdk1-PP2A signaling in controlling the orderly degradation of APC/C substrates, ensuring precisely timed assembly of the transcriptional infrastructure that coordinates cell cycle commitment and progression.

### Supplementary Figures and Figure legends

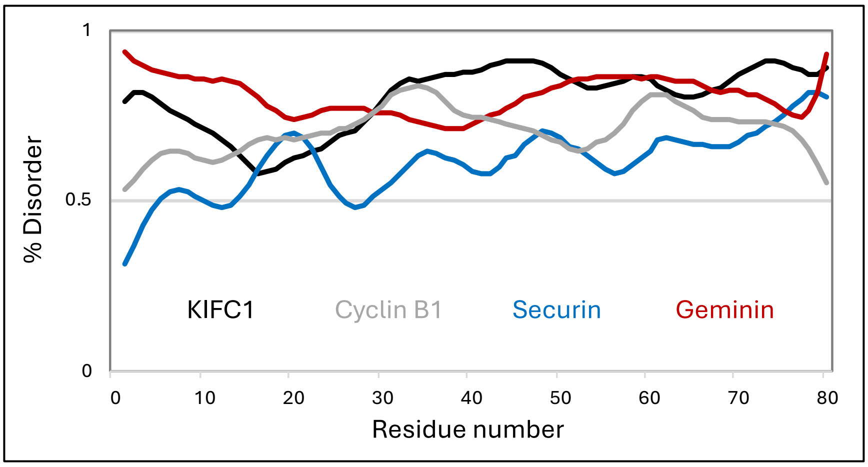


**Figure S1. Characterization of disorder in the N-terminal Regions of APC/C Substrates.** Sequence-based disorder predictions for the N-terminal regions of four canonical APC/C targets (depicted) were generated using IUPRED3, with the 50% disorder threshold highlighted. Degradation signals of all four proteins are located within the analyzed regions.

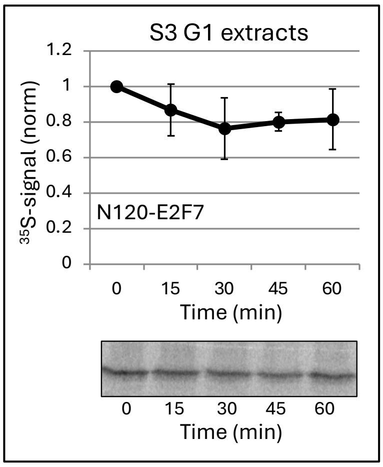


**Figure S2. N120-E2F7 is stable in G1 extracts.** Time-dependent degradation of E2F7 N-terminal fragment containing residues 1-120. Mean $^{35}$S signal and SEM are plotted; n = 3. Assays were analyzed by SDS-PAGE and autoradiography. Representative source data are shown.

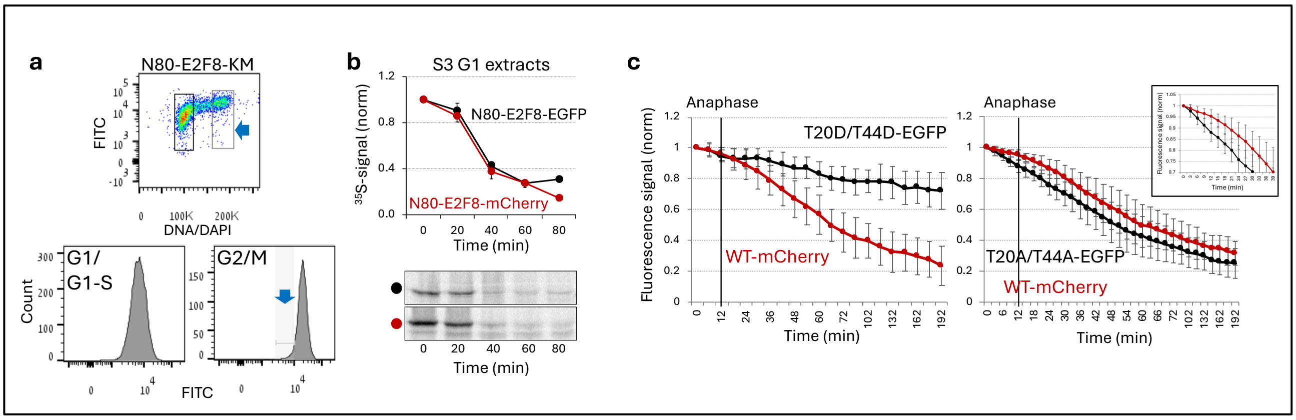


**Figure S3. Supplementary Information for Figure 3. (a)** Flow cytometry analysis of asynchronous HeLa cells expressing the KEN-box mutant (KM) N80-E2F8-EGFP. Top: Bivariate plots showing DNA content (DAPI) versus EGFP fluorescence, with G1 and G2/M phases highlighted (black and gray frames, respectively). Bottom: Histograms depicting EGFP signal distributions within these cell cycle phases. Blue arrows indicate G2/M subpopulations with relatively low EGFP levels. **(b)** EGFP/mCherry-tagged N80-E2F8 exhibits similar degradation kinetics. Time-dependent degradation of N80-E2F8-EGFP and N80-E2F8-mCherry (IVT, $^{35}$S-labeled) in S3 G1 extracts. Degradation assays were analyzed by SDS-PAGE and autoradiography. Top: Plot showing mean $^{35}$S signal and SEM; n = 3. Bottom: Color-coded representative source data. **(c)** Time-lapse microscopy of HeLa cells co-expressing N80-E2F8-mCherry and either T20D/T44D or T20A/T44A variants tagged with EGFP. Mean and SDM fluorescence intensities measured at 3- and 6-minute intervals are plotted (n = 20). Inset: Magnification of time-points 0-39 min. Anaphase is indicated by a vertical line. Equivalent analysis but with reciprocal tagging is presented in Figure 3f.

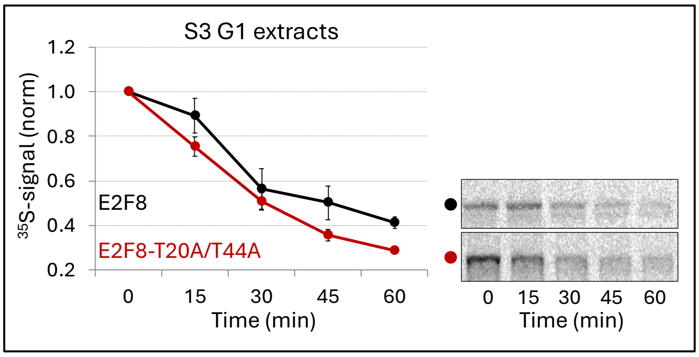


**Figure S4. Accelerated degradation of non-phosphorylatable E2F8 in G1 extracts.** Time-dependent degradation of WT- and Thr20/44-to-Ala E2F8 variants (IVT, $^{35}$S-labeled) in S3 G1 extracts. Protein degradation was assessed by SDS-PAGE and autoradiography. Mean $^{35}$S signal and SEM (n = 3) are plotted, with representative color-coded source data shown.

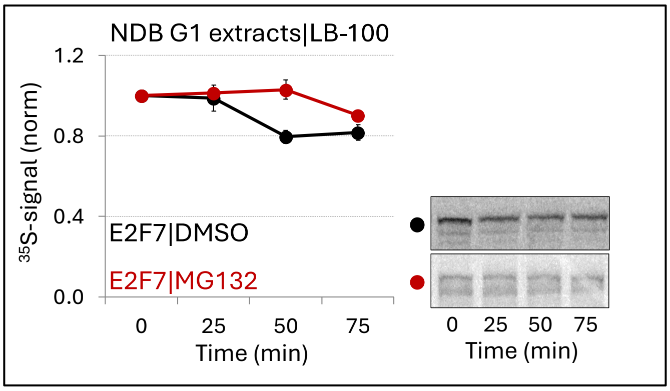


**Figure S5. Limited degradation of E2F7 in NDB G1-like extracts.** Time-dependent degradation of E2F7 ($^{35}$S-labeled IVT product) in NDB G1-like extracts treated with the PP2A inhibitor LB-100 and either the proteasome inhibitor MG132 or DMSO (control). Assays were analyzed by SDS-PAGE and autoradiography. Mean $^{35}$S signal intensity and SEM (n = 2-3) are plotted, with representative color-coded source data shown.

#### Table S1: A list of plasmids and cloning strategies

| Plasmid | New/ Lab stock | Donor plasmid | Acceptor plasmid | N-Tag | C-Tag | N-R.E | C-R.T | 5'-Primer/^3' Primer | Comments |
|---|---|---|---|---|---|---|---|---|---|
| *MBP-N80-E2F8-pET28 | New | His-N80-E2F8-EGFP pCS2-FA | MBP-WIP-pET28 | - | - | - | - | 5CTCCGCGGGTGAAAACCTGTACTTCCAGGGTGAGAACGAAAAGGAAAATCTCTTTTGTGAG3<br>5CTTTGTTTAGCAGCCTAGGTATTAATCAATTATTTCTGATCTCTGTTGCGGATCTCAGGGC3 | |
| *MBP-N100-E2F7-pET28 | New | His-N100-E2F7-EGFP pCS2-FA | MBP-WIP-pET28 | - | - | - | - | 5CTCCGCGGGTGAAAACCTGTACTTCCAGGGTGAGGTAAATTGTTTAACACTAAAAGACCTG3<br>5CTTTGTTTAGCAGCCTAGGTATTAATCAATTAGTCCCTTATATCTGGGCTGGCAGCACTAAT3 | |
| His-N80-E2F8-EGFP-pCS2-FA | Lab stock | | | | | | | | |
| His-N100-E2F7-EGFP-pCS2-FA | New | His-E2F7-pCS2-FA | His-N80-E2F8-EGFP-pCS2-FA | His | EGFP | FseI | AgeI | 5GCATGGCCGGCCCACCATGGAGGTAAATTGTTTAAC3<br>5ATGCACCGGTGGGTCCCTTATATCTGGGCTGGC3 | |
| His-N100-E2F7Δ-EGFP-pCS2-FA | New | E2F7-pCS2-FA | His-N80-E2F8-EGFP-pCS2-FA | His | EGFP | FseI | AgeI | 5GCATGGCCGGCCCACCATGCAAAAGGAAAATATATTTGTTGAT3<br>5ATGCACCGGTGGGTCCCTTATATCTGGGCTGGC3 | |
| **His-N100-E2F7+P-EGFP-pCS2-FA | New | His-N100-E2F7-EGFP-pCS2-FA | - | His | EGFP | - | - | 5AAGGAAAATATATTTGTTGATCCACGATCAAGGATGGCC3<br>5GGCCATCCTTGATCGTGGATCAACAAATATATTTTCCTT3 | Pro inserted between residues 37 and 38 |
| **His-N100-E2F7Δ+P-EGFP-pCS2-FA | New | *His-N100-E2F7Δ-EGFP-pCS2-FA | - | His | EGFP | - | - | 5AAGGAAAATATATTTGTTGATCCACGATCAAGGATGGCC3<br>5GGCCATCCTTGATCGTGGATCAACAAATATATTTTCCTT3 | Pro inserted between residues 37 and 38 |
| E2F7-pCS2-FA | Lab stock | | | | | | | | |
| E2F8-pCS2-FA | Lab stock | | | | | | | | |
| Securin-pCS2-FA | Lab stock | | | | | | | | |
| His-N80-E2F8-T20A/T44A-EGFP-pCS2-FA | Lab stock | | | | | | | | Thr20 and Thr44 are substituted with Ala |
| His-N80-E2F8-T20D/T44D-EGFP-pCS2-FA | Lab stock | | | | | | | | Thr20 and Thr44 are substituted with Asp |
| **His-N80-E2F8-T20A/T44A/KM-EGFP-pCS2-FA | New | His-N80-E2F8-T20A/T44A-EGFP-pCS2-FA | - | His | EGFP | | | 5GGCCCACCATGGAGAACGAAGCGGCAAATCTCTTTTGTGAGCCAC3<br>5GTGGCTCACAAAAGAGATTTGCCGCTTCGTTCTCCATGGTGGGCC3 | KEN at position 5-7 is substituted with AAN. Thr20 and Thr44 are substituted with Ala |
| E2F8 T20A/T44A-pCS2-FA | Lab stock | | | | | | | | Thr20 and Thr44 are substituted with Ala |
| E2F8 T20D/T44D-pCS2-FA | Lab stock | | | | | | | | Thr20 and Thr44 are substituted with Asp |
| **E2F8 T20A/T44A-KM-pCS2-FA | New | E2F8 T20A/T44A -pCS2-FA | - | - | - | - | - | 5GGCCCACCATGGAGAACGAAGCGGCAAATCTCTTTTGTGAGCCAC3<br>5GTGGCTCACAAAAGAGATTTGCCGCTTCGTTCTCCATGGTGGGCC3 | KEN at position 5-7 is substituted with AAN. Thr20 and Thr44 are substituted with Ala |
| **His-N100-E2F7-T45A- | New | His-N100-E2F7- | - | His | EGFP | - | - | 5CGATCAAGGATGGCCCCGAAGGCTCCAATAAAAAATGA3 | |

| | | | | | | | | | |
|---|---|---|---|---|---|---|---|---|---|
| EGFP-pCS2-FA | | EGFP-pCS2-FA | | | | | | 5TCATTTTTTATTGGAGCCTTCG GGGCCATCCTTGATCG3 | Thr45 is substituted with Ala |
| **His-N100-E2F7-T68A-EGFP-pCS2-FA | New | His-N100-E2F7-EGFP-pCS2-FA | - | His | EGFP | - | - | 5ACTCCAGAAAGAAATCCCATT GCTCCAGTTAAGCTTGT3<br>5ACAAGCTTAACTGGAGCAATG GGATTTCTTTCTGGAGT3 | Thr68 is substituted with Ala |
| **His-N100-E2F7-T45A/T68A-EGFP-pCS2-FA | New | His-N100-E2F7-T45A-EGFP-pCS2-FA | - | His | EGFP | - | - | 5ACTCCAGAAAGAAATCCCATT GCTCCAGTTAAGCTTGT3<br>5ACAAGCTTAACTGGAGCAATG GGATTTCTTTCTGGAGT3 | Thr45 and Thr68 are substituted with Ala |
| **E2F7 T45A/T68A-pCS2-FA | New | E2F7-pCS2-FA | - | - | - | - | - | 5CGATCAAGGATGGCCCCGAA GGCTCCAATAAAAAATGA3<br>5TCATTTTTTATTGGAGCCTTCG GGGCCATCCTTGATCG3<br>5ACTCCAGAAAGAAATCCCATT GCTCCAGTTAAGCTTGT3<br>5ACAAGCTTAACTGGAGCAATG GGATTTCTTTCTGGAGT3 | Thr45 and Thr68 are substituted with Ala |
| **E2F7 T45D/T68D-pCS2-FA | New | E2F7-pCS2-FA | - | - | - | - | - | 5CGATCAAGGATGGCCCCGAA GGATCCAATAAAAAATGA3<br>5TCATTTTTTATTGGATCCTTCG GGGCCATCCTTGATCG3<br>5ACTCCAGAAAGAAATCCCATT GATCCAGTTAAGCTTGT3<br>5ACAAGCTTAACTGGATCAATG GGATTTCTTTCTGGAGT3 | Thr45 and Thr68 are substituted with Asp |
| His-N80-E2F8-KM-EGFP-pCS2-FA | Lab stock | | | | | | | | |
| His-N80-E2F8-mCherry-pCS2-FA | New | pmCherry-N1 | His-N80-E2F8-EGFP-pCS2-FA | His | mCherry | AgeI | AscI | 5CCACCGGTCGCCACCATGG TGAGCAAG3<br>5GATGGCGCGCCCTACTTGTA CAGCTCGT3 | |
| His-N120-E2F7-EGFP-pCS2-FA | New | E2F7-pCS2-FA | His-N80-E2F8-EGFP-pCS2-FA | His | EGFP | FseI | AgeI | 5GCATGGCCGGCCCACCATG GAGGTAAATTGTTTAAC3<br>5ATGCACCGGTGGATCTGTAAA TGCATCGTCCTT3 | |
| N100-E2F7-EGFP-PTZV | New | His-N100-E2F7-EGFP-pCS2-FA | PTZV | - | EGFP | SpeI | NotI | 5GCGGACTAGTATGGAGGTAAA TTGTTTAAC3<br>5GCATGCGGCCGCTTACTTGT ACAGCTCGTCCA3 | |
| mCherry-PTZV | New | pmCherry-N1 | PTZV | - | - | SpeI | NotI | 5ATCCACTAGTCACCATGGTGA GCAAGGGCGAG3<br>5GCATGCGGCCGCCTACTTGT ACAGCTCGTCCA3 | |
| N80-E2F8-mCherry-PTZV | New | His-N80-E2F8-mCherry-pCS2-FA | PTZV | - | mCherry | SpeI | NotI | 5GCGGACTAGTATGGAGAACG AAAAGGAAAA3<br>5GCATGCGGCCGCCTACTTGT ACAGCTCGTCCA3 | |
| N80-E2F8 T20A/T44A-mCherry-PTZV | New | His-N80-E2F8-T20A /T44A-mCherry-pCS2-FA | PTZV | - | mCherry | SpeI | NotI | 5GCGGACTAGTATGGAGAACG AAAAGGAAAA3<br>5GCATGCGGCCGCCTACTTGT ACAGCTCGTCCA3 | |
| N80-E2F8 T20D/T44D-mCherry-PTZV | New | His-N80-E2F8-T20D /T44D-mCherry-pCS2-FA | PTZV | - | mCherry | SpeI | NotI | 5GCGGACTAGTATGGAGAACG AAAAGGAAAA3<br>5GCATGCGGCCGCCTACTTGT ACAGCTCGTCCA3 | |
| N80-E2F8-EGFP-PTZV | New | His-N80-E2F8-EGFP-pCS2-FA | PTZV | - | EGFP | SpeI | NotI | 5GCGGACTAGTATGGAGAACG AAAAGGAAAA3<br>5GCATGCGGCCGCTTACTTGT ACAGCTCGTCCA3 | |
| N80-E2F8-T20A/T44A-EGFP-PTZV | New | His-N80-E2F8-T20A/T44 A-EGFP-pCS2-FA | PTZV | - | EGFP | SpeI | NotI | 5GCGGACTAGTATGGAGAACG AAAAGGAAAA3<br>5GCATGCGGCCGCTTACTTGT ACAGCTCGTCCA3 | |
| N80-E2F8-T20D/T44D-EGFP-PTZV | New | His-N80-E2F8-T20D/T44 D-EGFP-pCS2-FA | PTZV | - | EGFP | SpeI | NotI | 5GCGGACTAGTATGGAGAACG AAAAGGAAAA3<br>5GCATGCGGCCGCTTACTTGT ACAGCTCGTCCA3 | |
| N80-E2F8-KM-EGFP-PTZV | New | His-N80-E2F8-KM-EGFP-pCS2-FA | PTZV | - | EGFP | SpeI | NotI | 5GCGGACTAGTATGGAGAACG AAGCGGCAAA3<br>5GCATGCGGCCGCttACTTGTA CAGCTCGTCCA3 | |

^3’-primers are presented as reverse complement

*Plasmids generated by assembly PCR

**Plasmids generated by site-directed mutagenesis (Agilent; #20521)
